# An alpine plant shows no decrease in genetic diversity associated with rapid post-glacial range expansion

**DOI:** 10.1101/2024.11.04.621891

**Authors:** Mackenzie Urquhart-Cronish, Colette S. Berg, Dylan Moxley, Olivia J. Rahn, Lila Fishman, Amy L. Angert

## Abstract

While range expansion is hypothesized to be a mechanism for species persistence under climate change, many eco-evolutionary models describe demographic and genetic processes during range expansion that may decrease genetic variation and increase genetic load at the leading edge (i.e., expansion load). These predictions are related to dispersal limitation at the leading edge driving colonization dynamics, a scenario common in post-glacial range expansion at the continental scale (∼20,000 years ago). However, post-glacial range expansion can also occur on contemporary time scales, such as alpine glacier recession following the end of The Little Ice Age (∼150 years ago) and our understanding of dispersal limitation structuring these instances of rapid range expansion are relatively understudied. Here, we test whether there is evidence supporting the role of dispersal limitation during range expansion following alpine glacier retreat using the native alpine plant *Erythranthe (Mimulus) lewisii* by examining patterns of neutral genetic diversity (single nucleotide polymorphisms) across the history of glacier recession (i.e., glacier chronosequence) across two glacier forelands in Garibaldi Provincial Park, BC. We find weak support for the prediction of increasing clines in genetic differentiation towards the range edge, and no support for decreasing clines in genetic diversity, suggesting dispersal limitation is not characterizing colonization during range expansion, with the implication that the accumulation of expansion load at the range edge is likely not applicable on these spatiotemporal scales. Together, our results suggest that loss of genetic diversity for range-shifting species in the alpine is likely not a key contributing factor to any decreased fitness over time.

## Introduction

Under changing climatic conditions, many species are shifting their distributions polewards and upslope to track their climatic niches (Chen et al., 2011; Parmesan and Yohe, 2003). While range expansion is hypothesized to be a mechanism for species persistence under climate change, many eco-evolutionary models describe demographic and genetic processes during range expansion that may decrease genetic variation and increase the frequency of deleterious mutations at the leading edge (i.e., allele surfing) (Klopfstein et al., 2006; Travis et al., 2007), undermining individual fitness and potentially population persistence (i.e., expansion load) (Peischl et al., 2013, 2015). The original theory of expansion load (Peischl et al., 2013) demonstrated that the greatest amount of genetic load is most likely to accumulate under conditions of limited dispersal across homogeneous environments. Following the initial model, more ecologically complex models of expansion load have been developed to include the evolution of dispersal ability (Peischl and Gilbert, 2020) and expansion along habitats with underlying environmental gradients (Gilbert et al., 2017).

These more ecologically complex models both demonstrated a reduction in the number of deleterious mutations accumulating during range expansion compared to Peischl et al. (2013), and an associated “rescue effect” from expansion load. In both studies, deleterious mutation accumulation and the resulting expansion load was reduced because founder events became less drastic, resulting in greater population densities at the range edge and a higher efficacy of selection at the expansion front. While these theoretical studies (Gilbert et al., 2018; Peischl et al., 2013; Peischl and Gilbert, 2020) make robust predictions about how dispersal limitation affects population size and genetic diversity during range expansion, it remains unclear when natural populations exist in the parameter space that does result in the accumulation of deleterious mutations and expressed genetic load at the range edge. By examining genetic signatures of dispersal limitation during range expansion, even before assessing genetic signatures of putatively deleterious mutations, we can begin to evaluate the relevance of these negative genetic consequences in natural populations.

Empirical phyloegoegraphic studies have demonstrated how patterns of genetic diversity change with post-glacial range expansion (Hewitt, 2000). A meta-analysis found that historical range dynamics related to post-glacial colonization consistently show higher genetic diversity in older, refugial populations relative to younger, recolonizing populations (73% of studies support claim) and are often associated with a latitudinal gradient in genetic diversity (Pironon et al., 2017). These patterns of decreasing genetic diversity and increasing genetic differentiation after post-glacial range expansions are consistent with dispersal limitation: colonization characterized by serial founder events and sequential population bottlenecks occurring at the leading edge. These underlying demographic processes suggest that the accumulation of expansion load should also be relevant under post-glacial range expansion scenarios. However, there is a bias towards studying post-glacial range expansion on long spatial and temporal scales (i.e., across latitude and Quaternary interglacial cycles) (Davis and Shaw, 2001; Hewitt, 2000; Kuchta and Tan, 2005; Lee-Yaw et al., 2008; Tiedemann et al., 2004). Post-glacial range expansion also occurs on contemporary spatiotemporal scales, such as following alpine glacier retreat since the end of the Little Ice Age (i.e., across elevation, over ∼150 years) (Naftz et al., 1996). Our understanding of patterns of dispersal limitation and its effects on genetic diversity at these smaller scales is relatively underdeveloped (Raffl et al., 2008). As such, it is currently unclear whether these instances of post-glacial range expansion are limited by the rate of dispersal at the leading edge (i.e., serial founder events leading to low range-edge densities) or the rate of ice retreat (i.e., with time to build up population density at the range edge before subsequent colonization events). If dispersal limited, then post-glacial range expansion on smaller spatiotemporal scales may also be subject to the accumulation of expansion load.

A handful of previous studies have investigated patterns of intraspecific genetic diversity and genetic differentiation following alpine glacier retreat (Pluess, 2011; Powolny et al., 2016; Raffl et al., 2008, 2006). These studies used previous knowledge of glacier recession from the historical glacial maxima (oldest terminal moraine) to the contemporary glacial extent (i.e., the glacier foreland) to apply a space-for-time substitution to follow the pathways of post-glacial colonization (i.e., chronosequence approach) (Walker et al., 2010). They sampled along existing chronosequences (i.e., a gradient of terrain age at a single site) to assess patterns of genetic differentiation and diversity along the pathway of range expansion. Together, they have described patterns contrary to those predicted by parameter space that supports the accumulation of expansion load (i.e., no evidence of dispersal limitation); studies have found increasing (Powolny et al., 2016) or no detected change in the development of genetic diversity within colonizing populations (Pluess, 2011; Raffl et al., 2008, 2006) and mixed support for increased genetic differentiation along the glacier foreland chronosequence zones (yes: Raffl et al. 2008, 2006, no: Pluess 2011; Powolny et al. 2016). Among these four case studies, focal species have represented a variety of life histories (e.g., long vs. short lifespan, wind vs. insect pollinated, capable or not capable of clonal growth) and foreland size (1 - 5 km). However, all studies to date were conducted in the European Alps and used older types of genetic markers (AFLPs and microsatellites) that generated relatively fewer polymorphic loci than modern sequencing methods (i.e., single nucleotide polymorpisms or SNPs). To further test the role of dispersal limitation during post-glacial range expansion on relatively shorter spatiotemporal scales, we sampled *Erythranthe (Mimulus) lewisii*— Northern pink monkeyflower—a perennial alpine plant that is an early colonizer of disturbed riparian habitats, along previously mapped alpine glacier recession pathways (Koch et al., 2009) in Garibaldi Provincial Park, BC, Canada. We used modern sequencing methods to generate a dataset with thousands of genetic markers (SNPs), resulting in a test with more statistical power to detect differences in genetic diversity than previous studies.

In this study, we refer to “isolation-by-chronodistance” to examine patterns of genetic differentiation incorporating space-for-time substitution and “isolation-by-topographic-distance” to refer to spread over space only. We hypothesize that dispersal limitation structures genetic diversity during range expansion along alpine glacier forelands, setting the stage for founder events, genetic bottlenecks, and expansion load. If dispersal is limited during range expansion, we predict genetic signatures of isolation-by-chronodistance and decreasing genetic diversity along chronosequence zones. Alternatively, if dispersal during range expansion is not limited, patterns of isolation-by-chronodistance and negative clines in genetic diversity would not be detected. The alternative pattern could arise if initial founder events are followed by the eventual build up of high densities at the range edge, swamping genetic signatures of previously restricted dispersal, or if dispersal (via seed sources outside of the foreland) or gene-flow (via pollinator movement) is high across all chronosequence zones. Ultimately, we are interested in the lasting signatures of range expansion in the alpine and how our results fit in with the handful of existing studies to begin to create consensus about the likelihood that expansion load is a risk for species undergoing range expansion across short spatiotemporal timescales.

## Methods

### Study system

*Erythranthe lewisii* (Phrymaceae) or Northern pink-monkeyflower is an herbaceous perennial plant that is native to subalpine and alpine riparian habitat throughout Northwestern North America. Aspects of the natural history of *E. lewisii* (e.g., riparian, early colonizer of disturbed habitat) tie it to the ecology of the glacier foreland, making it an ideal focal species to study postglacial range expansion. It is primarily bumblebee pollinated, but possesses hermaphroditic flowers and is capable of self-fertilization (both within and between flowers) and can reproduce clonally via rhizomes. Seed dispersal in monkeyflowers is thought to primarily occur via water (Waser et al., 1982), but dispersal upslope along the glacier foreland could potentially occur via wind and mammals (Vickery et al., 1986) and seed dispersing from source populations located on moraine slopes above/outside the glacier foreland. Here, we focus on *E. lewisii* in Garibaldi Provincial Park (British Columbia, Canada) within two alpine glacier forelands: Helm and Garibaldi/Lava (Figure 1A, B, C). In Garibaldi Provincial Park, *E. lewisii* plants are found growing along slow-moving alpine streams, rivers, and seeps. Their growing season begins with snow melt (mid-July) and continues until snow fall (early September).

### Field collections along glacier forelands

Fieldwork took place in Garibaldi Provincial Park (located in southwestern British Columbia) during August and September 2020 (Letter of Authorization: 98700-20). Here, we sampled *E. lewisii* along two distinct alpine glacier forelands located (Fig. 1A) ∼18 km apart, accounting for topography, that have known timing of glacier retreat (Koch et al., 2009) (Fig. S1) where Koch et al. (2009) characterized the timing of glacier retreat using a combination of historical images, dendrochronology, and physical evidence of terminal and lateral moraines. We designed our field collections based on these mapped chronosequence zones (Helm: 1928 - 2003, ∼3 km; Garibaldi/Lava: 1850 - 1977, ∼5 km) (Fig. S1), including sites within and outside of the alpine glacier foreland where we found *E. lewisii* growing (i.e., in suitable habitat) by the border of chronosequence zones (Fig. 1BC) (Table 1). Per site, we sampled leaf tissue from ∼10-20 plants with at least ∼1 m distance between sampled plants to avoid sampling clonal genets and stored field-collected leaf tissue in manilla coin envelopes placed inside bags filled with non-toxic indicator desiccant (Dry & Dry, B0725LNZ24).

**Figure 1:**
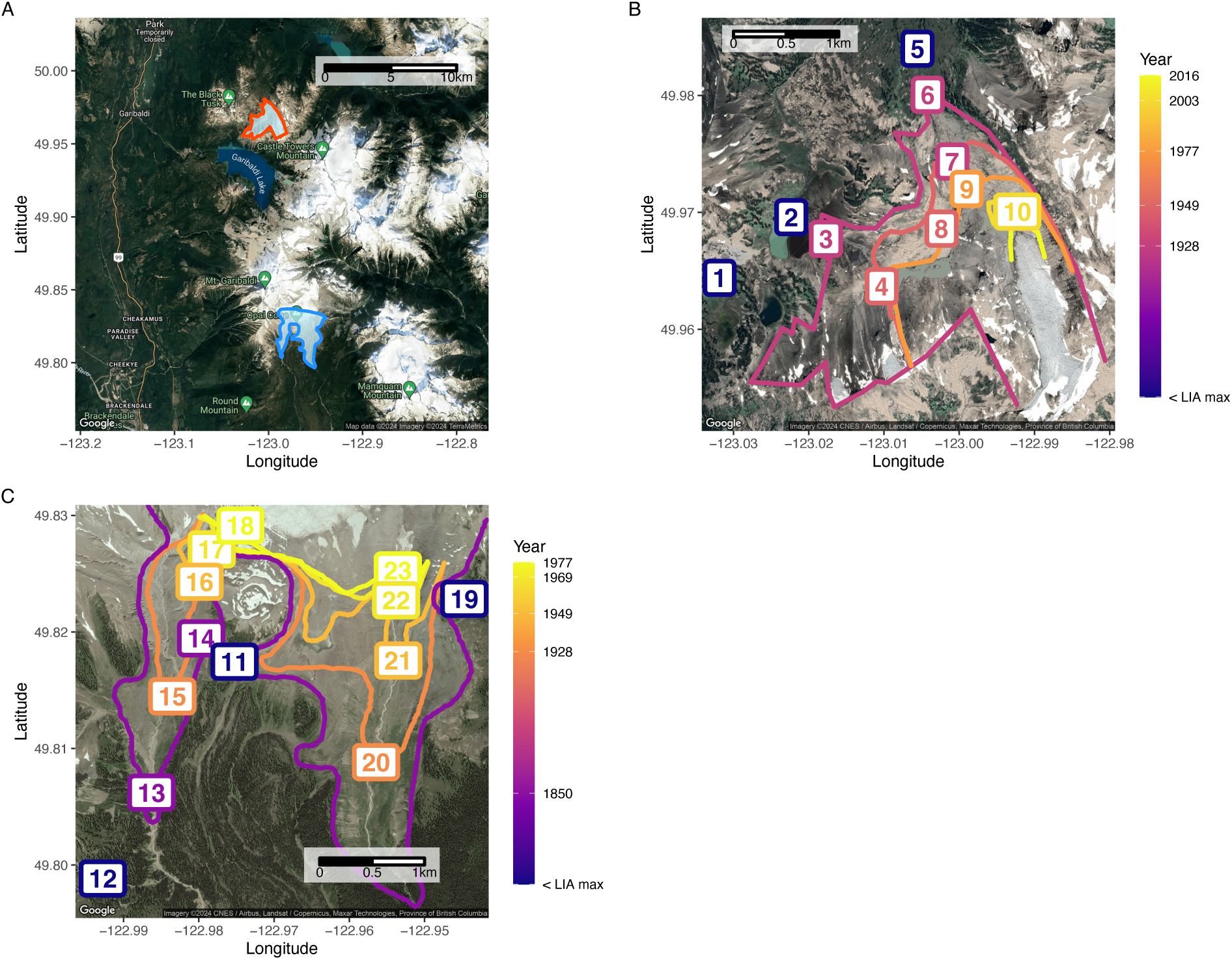
Study sites in Garibaldi Provincial Park, BC and Mapped glacier recession pathways with sampling locations. **A** Location of glacier forelands, with Little Ice Age (LIA) maximum mapped (Orange - Helm; Light blue - Garibaldi/Lava), sampled in this study. **B & C** Glacier recessions pathways mapped from Koch et al. (2009) and sampling locations for the B) Helm Glacier and C) Garibaldi/Lava Glacier forelands. Sampling locations are numbered by foreland (1 - 10 for the Helm foreland; 11 - 23 for the Garibaldi/Lava foreland) and year.

**Table 1:**
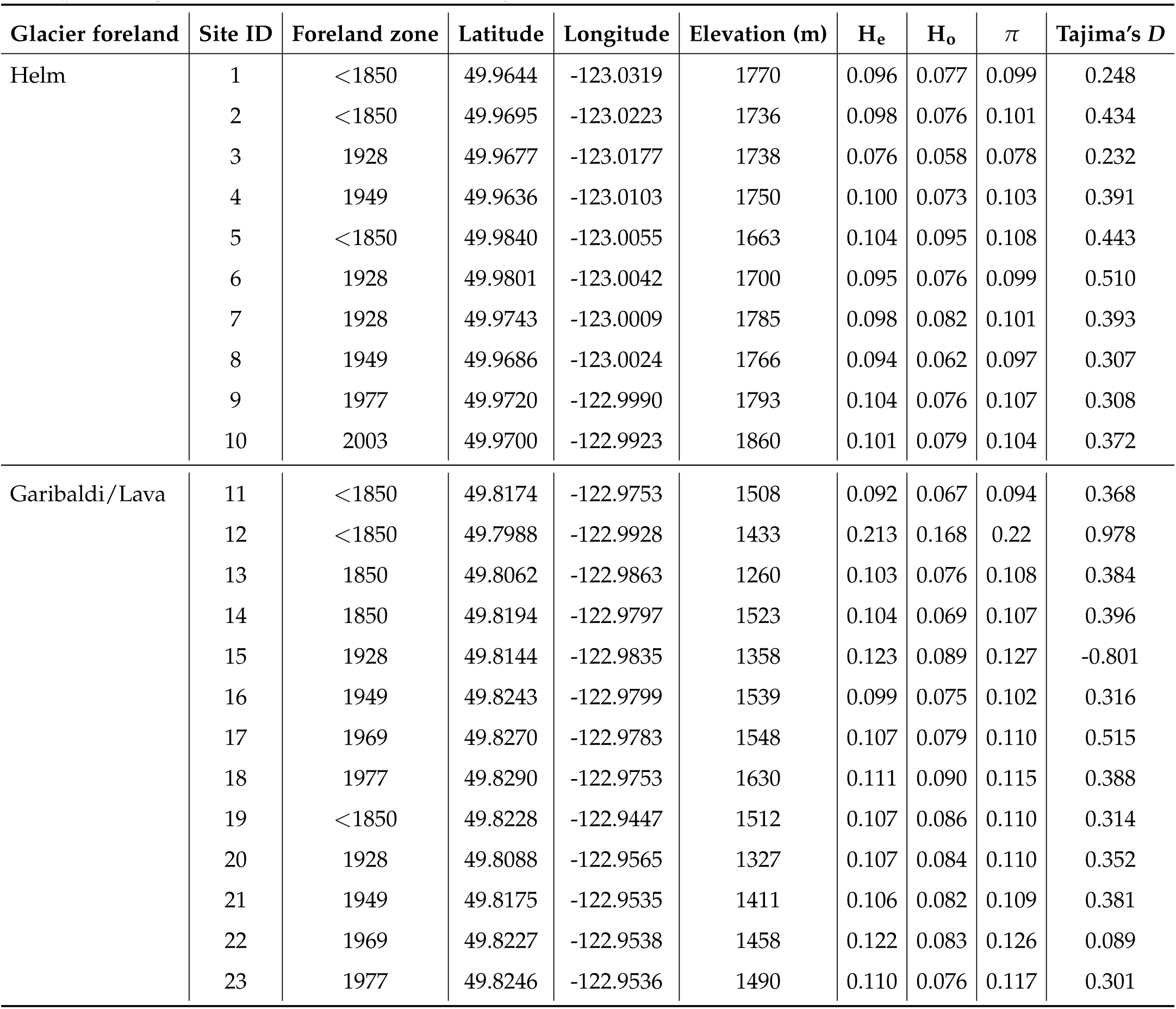
Locations of sampling sites and timing of glacier recession history within forelands with per-site genetic and nucleotide diversity data.

### ddRADseq genotyping

We extracted DNA from field-collected leaf tissue using a CTAB-chloroform protocol (full length protocol can be found at protocols.io) modified from Xin and Chen (2012). Briefly, we added 100 ug/mL Proteinase K (NEB P8107S) to the CTAB Lysis buffer to reduce enzymatic oxidation and increased centrifugation speeds to 6000 rcf following DNA-CTAB complex precipitation. For DNA purification, we suspended 5 uL of MagAttract beads (Qiagen 1026901) in 55 µL of DNA plus TE Buffer (10 mM TRIS, 0.1 mM EDTA, pH 8.0) and performed multiple washes. We transferred 54 µL of the sample to a clean PCR plate and sample purity was assessed using 2 µL of DNA elution on a Nanodrop 2000 (Thermo Scientific™), and DNA concentration was quantified with the Qubit 2.0 Broad Spectrum kit (Invitogen™) using 2 µL of DNA elution.

We then used a double-digest restriction-site associated DNA sequencing (ddRADseq) protocol to generate genome-wide sequence clusters (tags) for 357 individuals across 23 collection sites, following the BestRAD library preparation protocol) and methods from Kolis et al. (2022), initially using restriction enzymes *PstI* and *BfaI* (New England Biolabs, Ipswich, MA; both methylation insensitive) to digest DNA samples. After enzymatic digestion, sets of 48 individual DNA samples were labeled with unique in-line barcoded oligos, and each set pooled into a single tube. Pools were barcoded using NEBNext indexing oligos with a degenerate barcode, PCR amplified, and size selected (300 - 600 bp fragments) using a Bluepippin (Sage Science™) agarose gel. Libraries were prepared using NEBNext Ultra II library preparation kits for Illumina (New England Biolabs, Ipswich, MA). The libraries were paired-end sequenced (150 bp) on ^1^ lane of Illumina NovaSeq SP sequencer (University of Oregon Genomics and Cell Characterization Core Facility).

Illumina reads were demultiplexed and true PCR duplicates (same restriction site, plus same degenerate barcode) removed using a custom Python script. Adapters were trimmed and low-quality reads were removed using STACKS function process shortreads (Catchen et al., 2013, 2011).. Reads were mapped to the *E. erubescens* version2 reference genome (http://mimubase.org/FTP/Genomes/LF10g_v2.0/) using BWA MEM (Li and Durbin, 2009) and indexed with SAMtools (Li et al., 2009). SNPs were called using GATK (McKenna et al., 2010) for aligned sequence data. After alignment and quality filtering, we retained 102,835 sites (quality > 30, maximum missing data = 0.2). After filtering for sites with mean depth (≥10× & ≤30×), and mean depth per individual ≥7×, we retained 64316 variable sites for 332 individuals (Helm N=147 and Garibaldi/Lava N=185) representing all 23 collection sites (8 - 17 samples per site). For analyses of genetic diversity, we used genomic data where we did not filter for minor allele frequencies or linkage disequilibrium. For analyses of population structure we filtered minor allele frequency (maf) > 0.1 with bcftools (Danecek et al., 2021) and linkage disequilibrium with PLINK (Purcell et al., 2007) and retained a total of 6, 358 variable sites.

### Analyses of population genetic structure, differentiation, and diversity

#### Population genetic structure & genetic differentiation

To analyze population genetic structure we generated a dataset for principal component analysis using PLINK (Purcell et al., 2007) to summarize the overall genetic similarity of individuals and ran the program ADMIXTURE (Alexander et al., 2009) for *K* = 1 − 23 (maximum number of sites in sampling) to generate individual-level ancestry assignments. For principal component analysis (PCA), we calculated the 95% confidence levels of a-priori foreland assignment using the stat elipse() function from the *ggplot2* package assuming a multivariate t-distribution since the data were not normally distributed.

We calculated pairwise genetic distances (Weir and Cockerham’s *F_ST_*) between forelands and all sampled sites using functions from the R package *heirfstat* (Goudet and Jombart, 2022) within the R package *graph4lg* (Savary et al., 2021). We generated two measures of geographic distance, *1)* represents the space-for-time substitution of the chronosequence approach where we calculated pairwise differences in year of glacier recession zone to represent the pathway of range expansion and *2)* “isolation-by-topographic-distance” represents topographic distances between sampling sites (without the spatiotemporal context of range expansion). To calculate topographic distances we accessed topographic data via the open source website OpenTopography and used the R package *elevatr* (Hollister et al., 2023) to generate a topographic raster layer of the area of our study sites. We then used the R package *topoDistance* (Wang, 2020) to calculate a pairwise matrix of topographic distances between all sampled sites. To test for patterns of isolation-by-chronodistance and isolation-by-topographic-distance within distinct forelands, we used the R packages *ade4* (Dray and Dufour, 2007) and *vegan* (Oksanen et al., 2022) to run separate Mantel tests per site, with 9999 permutations.

#### Genetic diversity

We used the R package *graph4lg* to calculate the following measures of genetic diversity within each distinct site: mean observed heterozygosity (*H_o_*), mean expected heterozygosity (*H_e_*), and mean allelic richness. We tested the relationship between chronosequence zone, foreland identity, and genetic diversity by running a linear model using the R package *lme4* (Bates et al., 2015) to test the individual relationship and interactions between time (chronosequence zone) and genetic diversity across forelands.

To investigate nucleotide diversity, we used an “all sites” vcf (including variant and invariant sites) to avoid biasing our estimates of nucleotide diversity (Korunes and Samuk, 2021) generated with GATK (McKenna et al., 2010) and filtered for depth and missing data in the same way as above (retained 96,795 sites). We then used the software *pixy* (Korunes and Samuk, 2021) to calculate average nucleotide diversity within (*π*) populations over a 1 Mb window per chromosome. We then averaged the per-region values to generate a per individual value.

We calculated average values of Tajima’s *D*—a method to detect shifts in the site frequency spectrum away from neutral expectations by comparing the number of segregating sites and average pairwise base pair differences—-using vcftools (Danecek et al., 2011) in a similar way, using our “all sites” vcf with a 1 Mb window size and averaging values per site and then per population. Tajima’s *D* is a useful statistic to make inferences whether deviations from neutral evolutionary expectations are occurring, due to selection and/or demography. When values are strongly positive that suggests an excess of intermediate-frequency variation in observed samples and this can be driven demographically by a recent population bottleneck or as a result of balancing selection; if values are strongly negative, this suggests a deficit of observed genetic variation/an excess of singletons, and this can be driven demographically by recent population growth or as a result of positive selection/selective sweeps (Hahn, 2019). We were interested in making inferences about demography (recent founder events) and tested the relationship between chronosequence zone and Tajima’s *D*. We ran a linear model using the R package *lme4* to test the relationship between space/time (interaction between numerical chronosequnce zones and categorical forelands) and deviations from neutral evolution.

## Results

### Distinct population structure detected between and within forelands

In the principal component analysis, PC1 explained 15.6% and PC2 explained 10.1% of variation in population genetic diversity (Fig. 2A). Individuals from each foreland clustered in distinct groups along PC1, as demonstrated by ellipses depicting the 95% confidence levels of foreland identity. We also found evidence of weak genetic differentiation between forelands (*F_ST_* = 0.035). While PC1 separated individuals between foreland sites, PC2 appeared to summarize genetic structure present within sites, specifically identifying genetic dissimilarity between a single site within the Helm foreland (site 3, Fig. S2A) and elevated genetic differentiation relative the rest of the sampled sites within the foreland (Fig. S3).

Based on our sampling design (two forelands) and the visual representation of genetic structure defined by the PCA (two to three distinct clusters) we plotted the ADMIXTURE results for *K* = 2 (Fig.2B) and *K* = 3 (Fig. S2B) ancestral clusters. We also examined the CV error plot from the ADMIXTURE analysis, however; clustering algorithms can overfit data easily and caution is recommended when interpreting these inferred values of K (Pritchard et al., 2000) (in our case, *K* = 7, 6, &8). For these reasons we based our ancestral cluster assignment on the visual structure in PC space (Fig.2A). Most individuals within sites were assigned mixed ancestry, but with the majority of ancestry assignment belonging to the respective foreland with which each individual was sampled. Site 3 in the Helm foreland, which had some individuals forming a distinct cluster along PC2 (site 3, Fig. S2A), were assigned to a distinct ancestry cluster when *K* = 3 (Fig. S2B).

**Figure 2:**
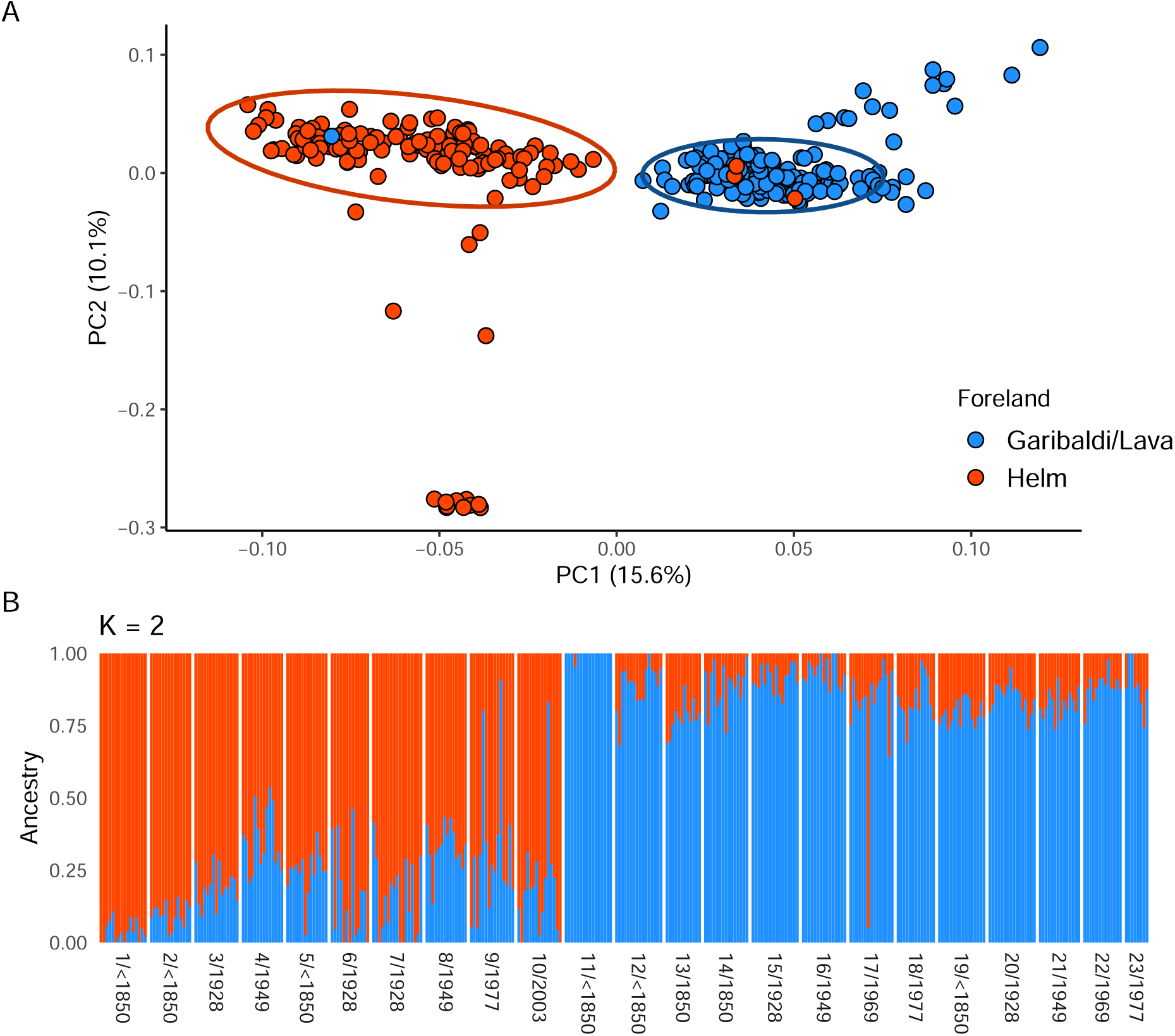
Population genetic structure suggests two distinct genetic groups with greater similarity among individuals sampled within a given foreland. A) PCA plot with data points coloured by sampling location (foreland) with ellipses representing the 95% confidence intervals around foreland identities (Orange = Helm, Blue = Garibaldi/Lava). B) ADMIXTURE analysis results for *K* = 2 plotting proportion assigned ancestry. X-axis labels the site (Fig. 1 BC) and the glacier recession zone; sites 1 - 10 are Helm and 11 - 23 are Garibaldi/Lava. Colours match the legend in panel A.

### Mixed evidence for isolation-by-chronodistance within forelands

To assess whether dispersal was limited across the foreland, we examined whether genetic differentiation increased with increasing distance from the historical range core to the contemporary range edge using pairwise comparisons of *F_ST_* and timing of glacier recession (our proxy for recency of range expansion). We found no support for isolation-by-chronodistance in the Helm foreland (*r* = −0.15, *p >* 0.05) (Fig. 3A) and a significant relationship in the Garibaldi/Lava fore-land (*r* = 0.39, *p* = 0.008) (Fig. 3B). We also investigated the relationship between topographic distance, ignoring the influence of range expansion history, and genetic distance and found no evidence for isolation-by-topographic-distance in either foreland (Fig. S4).

**Figure 3:**
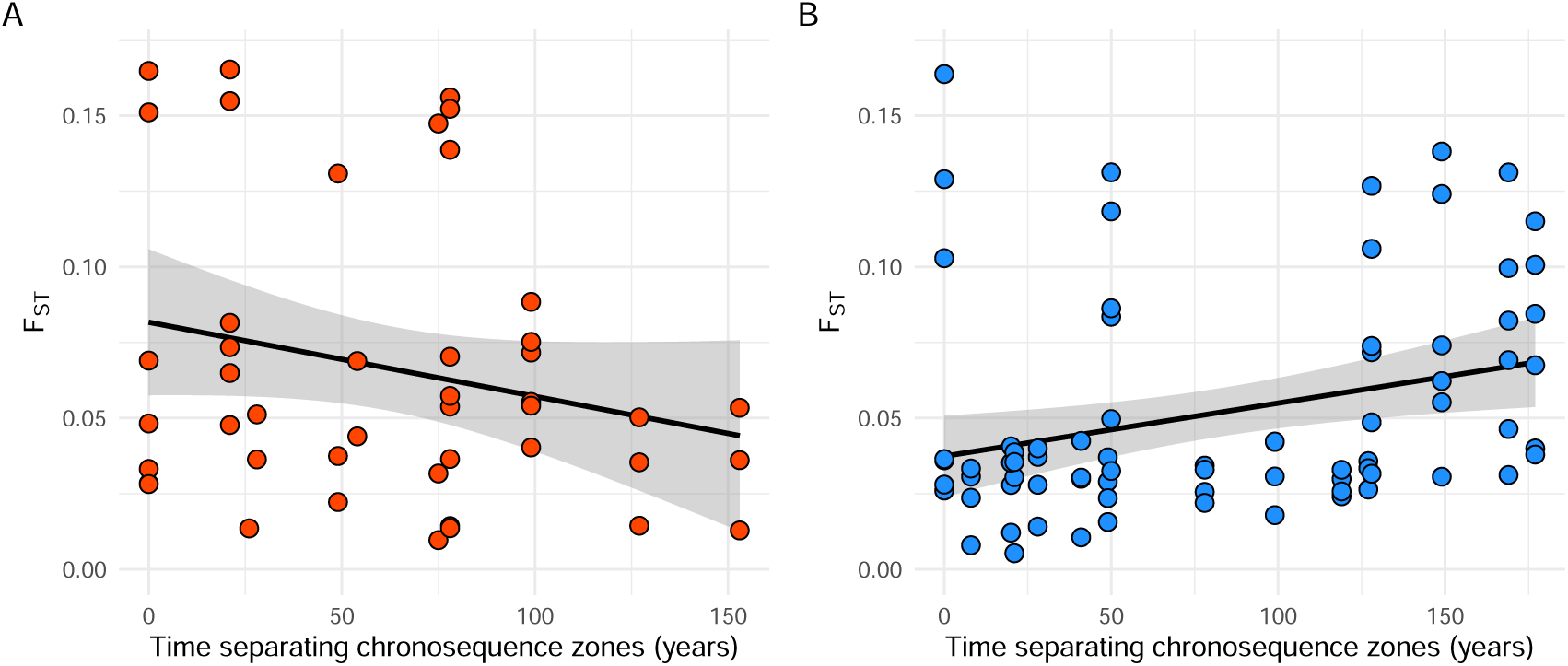
Relationships between range expansion history (years) versus genetic differentiation (*F_ST_*), evidence of restricted gene flow between glacier recession zones, within forelands are varied. **A)** Helm **B)** Garibaldi/Lava. Mantel tests with 9999 permutations are non significant for Helm (Mantel statistic *r* : −0.15, *p* = 0.79) and significant for Garibaldi/Lava (Mantel statistic *r* : 0.39, *p* = 0.0086). Linear regression was used to fit lines to the scatterplots.

### No clines in genetic diversity detected along pathways of range expansion

We examined patterns of genetic diversity following the timing of glacier retreat to detect patterns associated with founder effects (i.e., decreased heterozygosity) due to leading edge colonization, characteristic of limited dispersal during range expansion. We quantified *H_o_*, *H_e_*, and allelic richness for all sites sampled within forelands and used linear models to test the relationship between timing of expansion and levels of genetic diversity. Our model results revealed no relationship between timing of expansion and genetic diversity, foreland identity, or their interactions (Table 2). Comparisons between *H_e_* and *H_o_* within sites all showed *H_e_ > H_o_*, suggesting non-random mating, perhaps due to inbreeding, within sites. Tajima’s *D* values did not provide support for demographic signatures of recent population bottlenecks (or recent population growth) related to range expansion following alpine glacier retreat (chronosequence zone year: *β* = −0.03, SE = 0.05, *p* = 0.5; foreland: *β* = −0.03, SE = 0.3, *p* = 0.9; chronosequence zone year * foreland: *β* = 0.03, SE = 0.08, *p* = 0.7).

**Figure 4:**
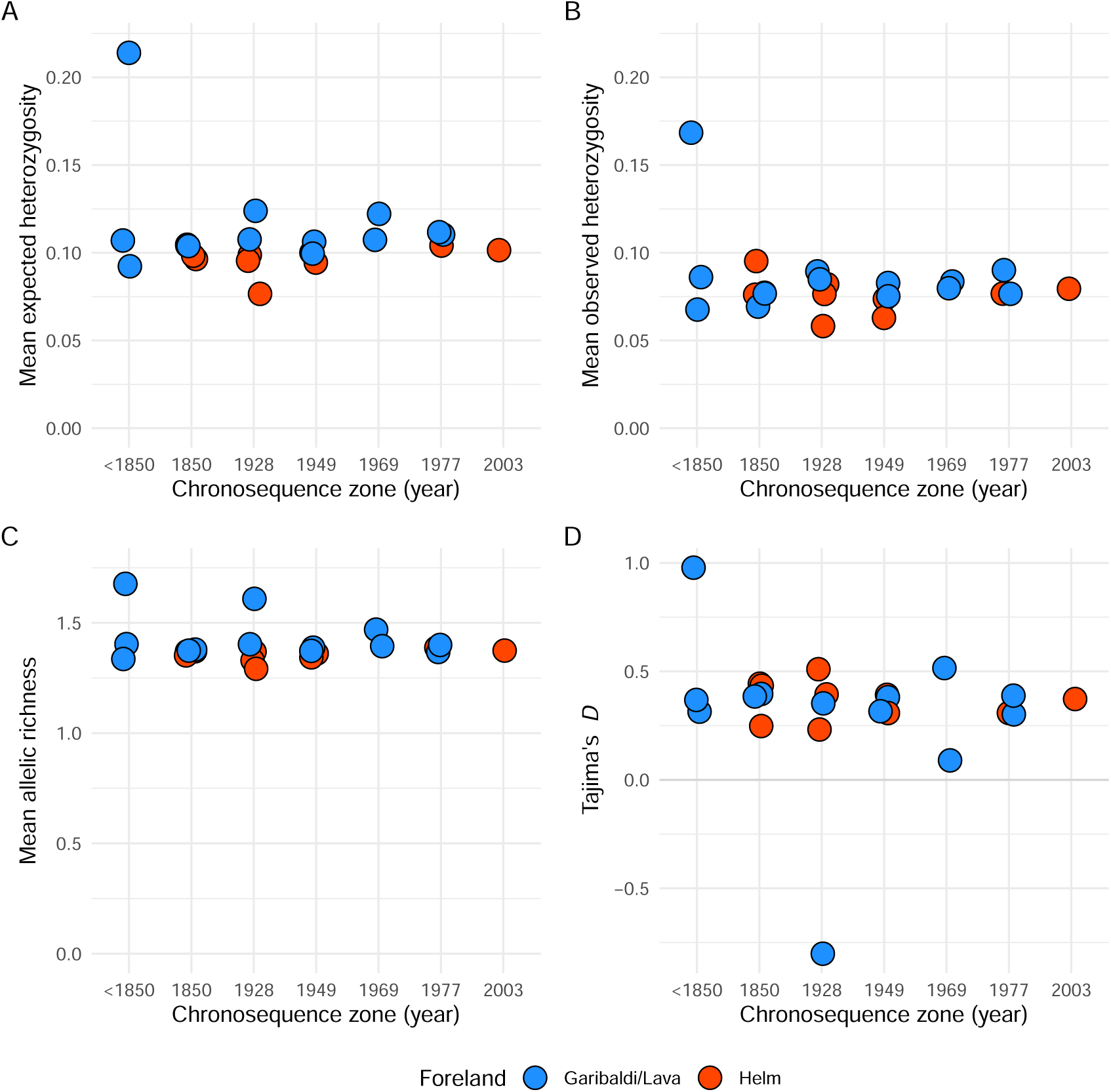
No evidence of population bottlenecks characterizing range expansion as measured by genetic (A, B, C) and nucleotide (D) diversity across chronosequence zones in each distinct foreland (Helm and Garibaldi/Lava) **A)** Mean expected heterozygosity by locus by site **B)** Mean observed heterozygosity by locus by site **C)** Mean allelic richness by locus by site, and **D)** Mean Tajima’s *D*, a metric that represents deviations from neutral evolutionary processes (demographic processes or selection). No relationships between timing of glacier recession and genetic diversity (A, B, C) or nucleotide diversity were detected. Note, due to differences in timing of glacier recession (Fig. S1), populations from Helm are absent from the < 1850 core zone and the 1850 time period represents habitat outside the LIA glacial maximum.

**Table 2:** Linear model results testing relationship between chronosequence zone years, fore-land identity, and their interactions on values of genetic diversity.

| Response variable | Variable | $\beta$ | SE | t value | P-value |
| --- | --- | --- | --- | --- | --- |
| $H_e$ | chronosequence zone year | -1.207e-04 | 9.549e-05 | -1.26 | 0.2 |
|  | foreland | -2.524e-01 | 3.347e-01 | -0.75 | 0.5 |
|  | chronosequence zone year * foreland | 1.225e-04 | 1.746e-04 | 0.7 | 0.5 |
| $H_o$ | chronosequence zone year | -1.085e-04 | 8.125e-05 | -1.3 | 0.2 |
|  | foreland | -9.925e-02 | 2.848e-01 | -0.35 | 0.7 |
|  | chronosequence zone year * foreland | 4.685e-05 | 1.486e-04 | 0.3 | 0.8 |
| Allelic richness | chronosequence zone year | -0.0002 | 0.0003 | -0.7 | 0.5 |
|  | foreland | -0.6 | 1.1 | -0.5 | 0.6 |
|  | chronosequence zone year * foreland | 0.0003 | 0.0006 | 0.4 | 0.7 |

## Discussion

In this study, we demonstrated that an alpine plant shows inconsistent evidence of isolation-by-chronodistance and no evidence of decreased genetic diversity during rapid post-glacial range expansion. In one of our two replicate forelands, we found that glacier recession zones that were farther apart in time had increased genetic differentiation, suggesting reduced gene flow between populations in older vs. more recently exposed habitat. However, we did not find genetic patterns consistent with strong or lasting dispersal limitation during expansion, suggesting either that founder effects during expansion were weak or any effects on genetic diversity were temporary. Overall, our results—combined with the existing literature—suggest that the negative genetic consequences predicted by the original theory of expansion load (Peischl et al., 2013) are not likely to be applicable to post-glacial range expansions on short spatiotemporal scales.

### Population structure emerging on small space and time scales

We detected evidence of population genetic structure between the two glacier forelands, even with only 18 km of topographic distance separating both sites. Genetic differentiation across this relatively small spatial scale could be due to landscape features that separate the two forelands, such as Garibaldi Lake and the Garibaldi névé (snow field and mountains) (Fig. 1A), that may prevent successful dispersal of seed and/or pollen between forelands. Other studies that have found fine-scale population genetic differentiation across topographically complex areas have attributed these patterns to local geographic features (e.g., mountain ridges, lakes, etc.) (Hotaling et al., 2018; Trense et al., 2021). We also detected potential evidence of isolation-by-environment (as opposed to a process related to range expansion) for a single site within the Helm foreland (Fig. S2A, B). Since site 3 belongs to an earlier chronosequence zone and not the range edge, this differentiation could be related to distinct ecological conditions (e.g., wetter, cooler climate) of the site, however we did not not quantify site-level conditions.

### Evaluating patterns of limited dispersal

#### Genetic differentiation

Within forelands, we found support for a pattern of isolation-by-chronodistance only in the Garibaldi/Lava foreland. Again, landscape differences at the scale of the individual foreland such as size, length of glacier recession history, and elevation (Table 1) may potentially underlie differences in degree of isolation-by-chronodistance at this relatively small spatial scale. For example, compared to the Helm foreland the Garibaldi/Lava foreland is larger, has longer recession history, and also has a larger elevational range, and these factors may aid in restricting gene flow among chronosequence zones following colonization with dispersal limitation. However, while the Helm foreland is approximately half the size of the Garibaldi/Lava foreland, previous work investigating patterns of intraspecific genetic variation in an alpine plant (*Trifolium pallescens*) along glacier recession pathways in Switzerland found significant patterns of isolation-by-chronodistance in a foreland as small as 1 km (Raffl et al., 2008). The riparian natural history of *E. lewisii*, specifically seed dispersal via water (i.e., downstream), could facilitate propagule mixing among younger and older chronosequence zones, decreasing overall levels of genetic differentiation, especially in the smaller Helm foreland.

Additionally, phenological differences in *E. lewisii* across the forelands may interact with foreland size in shaping spatial patterns of genetic differentiation. For example, differences across elevation in the timing of snow melt early in the season are expected to stagger phenological overlap within a foreland, influencing early season gene-flow dynamics (via pollinator activity) (Albrecht et al., 2010), and larger foreland size—such as the Garibaldi/Lava foreland relative to the Helm foreland—could further exacerbate these phenological limits to gene flow and preserve spatial genetic structure characterized by dispersal limitation during colonization. Differences in the timing of glacier recession may also matter, as supported by a meta-analysis that found weak evidence of an association between longer timescales of colonization history and positive signals of isolation-by-distance (Crispo and Hendry, 2005).

We evaluated patterns of dispersal-limitation by investigating spatial patterns in population genetic summary statistics. Other methods can be used to infer demographic pathways of range expansion, such as demographic inference using the site-frequency-spectrum and Approximate Bayesian Computation modelling (Marchi et al., 2021). We selected summary statistics since we believed the spatiotemporal scale of our study may have limited our use of more complex methods, but future work could assess the validity of these methods in detecting demographic change during range expansion in alpine glacier forelands (Epps and Keyghobadi, 2015).

#### Genetic diversity

Allelic and nucleotide diversity patterns support the alternative hypothesis that range expansion following alpine glacier retreat was buffered from the effects of genetic drift at the leading edge. The patterns we detected could be due to the absence of dispersal limitation during range expansion, or could arise from an initial instance of dispersal-limited during colonization that is transient. In the case of the absence of dispersal-limitation, the rate of expansion towards the leading edge may be determined by the rate of ice retreat rather than the rate of propagule movement, such that edge populations are able to reach high densities (conferring higher genetic diversity) before new habitat opens up via glacier recession for subsequent colonization. Alternatively, genetic signals of dispersal limitation could be obscured over time due to increased population densities at the range edge or if pollinator-facilitated gene flow increased levels of genetic diversity across chronosequence zones post-range expansion. While these different processes (i.e., no dispersal limitation vs. transient dispersal limitation) would lead to the same pattern of genetic diversity, attributing pattern to process would require additional work such as a pedigree study in the field to investigate the role of gene flow post-expansion (Gaudeul and Till-Bottraud, 2008).

### Genetic consequences of range expansion following alpine glacier recession

Based on our results, we can infer that the negative genetic consequences predicted by the original parameter space of models of expansion load are unlikely applicable on these spatiotemporal scales. This suggests range expansions following alpine glacier retreat are likely to have more in common with more recent ecologically-complex models of expansion load, allowing range edge populations to overcome the demographic effects of dispersal limitation (Gilbert et al., 2017; Peischl and Gilbert, 2020). We can extend our findings on genetic diversity to suggest plant populations undergoing range shifts in alpine ecosystems may be more resilient to ecological change than an influential and recent range expansion model suggests. Our research, combined with other studies on patterns of intraspecific genetic diversity in the alpine, shows range-edge populations do not experience decreased levels of neutral genetic diversity relative to the range core. Populations that are able to maintain levels of standing genetic variation during range expansion may possess greater evolutionary potential in the context of changing ecological conditions. More generally, our results lead to the inference that range expansions upslope and across elevation appear to experience different dispersal dynamics than their latitudinal counterparts, so researchers looking to apply theoretical models to distinct spatial scales in nature should be aware of the potential mismatch in predicted demographic pathways of range expansion.

### Conclusions

Our empirical study of post-glacial range expansion along alpine glacier recession pathways found weak support for the prediction of increasing clines in genetic differentiation and no support for decreasing clines in genetic diversity. While our results had greater statistical power to detect differences among genetic diversity markers than any previous studies, our results generally agree with previous studies that did not detect patterns of intraspecific genetic diversity on alpine glacier forelands. We did not find evidence in support of strong or lasting founder effects characterizing colonization during range expansion, with the implication that the accumulation of expansion load at the range edge is not applicable on these spatiotemporal scales. Together, our results suggest that loss of genetic diversity for range shifting species in the alpine is likely not contributing to their decreased fitness, and ecological interactions (i.e., novel competitors, herbivores, and pollinators) might be more important in driving patterns of species persistence in the face of climatic change (Richman et al., 2020; Speed et al., 2012; Usinowicz and Levine, 2021). Finally, we emphasize the importance of evaluating the essential underlying theoretical assumptions of model parameter space before extending investigation to conduct empirical tests of more complex eco-evolutionary theory in nature.

## Acknowledgements

Thanks to members of the Angert Lab, Loren Rieseberg, Jennifer Williams, Mike Whitlock, and Sally Aitken for helpful comments on earlier drafts of the manuscript. Thanks to Tom Booker and Julia Kreiner for assistance with genomic analyses. Thanks to Mannfred Boehem, Tom Booker, and Kelley Slimon for assistance with fieldwork and Garibaldi Provincial Park (Geoff Popowich & David Whiteside) for providing a Letter of Authorization to conduct fieldwork. Thanks to Tim Wheeler, David Xing, and the University of Montana Genomics Core for providing reagents, equipment, and support to complete ddRADseq library preparation. This research was enabled in part by support provided by the Digital Research Alliance of Canada (https://alliancecan.ca/en) .

## Data accessibility

### Data Accessibility Statement

All data and analysis code used in this article will be deposited in Zenodo upon publication. Sequencing data will be deposited in the NCBI Sequence Read Archive.

### Benefit-Sharing Statement

Benefits Generated: Benefits from this research accrue from the sharing of our data and results on public databases as described above.

## Author contributions

MUC and AA conceived of the study and designed the field sampling. MUC and OR conducted fieldwork. MUC conducted all DNA extractions with with training and assistance DM. MUC and CB conducted library prep lab work with LF providing space, reagents and assistance troubleshooting protocols. MUC ran the genomic analyses with demultiplexing scripts provided by CB. MUC generated and analyzed all data. MUC wrote the first draft of the manuscript with input and contributions from all authors. Funding from NSERC DG to ALA [RGPIN-2022-03113] and MUC (CGS-D graduate scholarship to MUC), the University of British Columbia (Four-year Fellowship and Tuition funds to MUC), and research grant funding from the Society for the Study of Evolution (SSE) R.C. Lewontin Early Award to MUC.

## Supplementary figures

**Figure S1:**
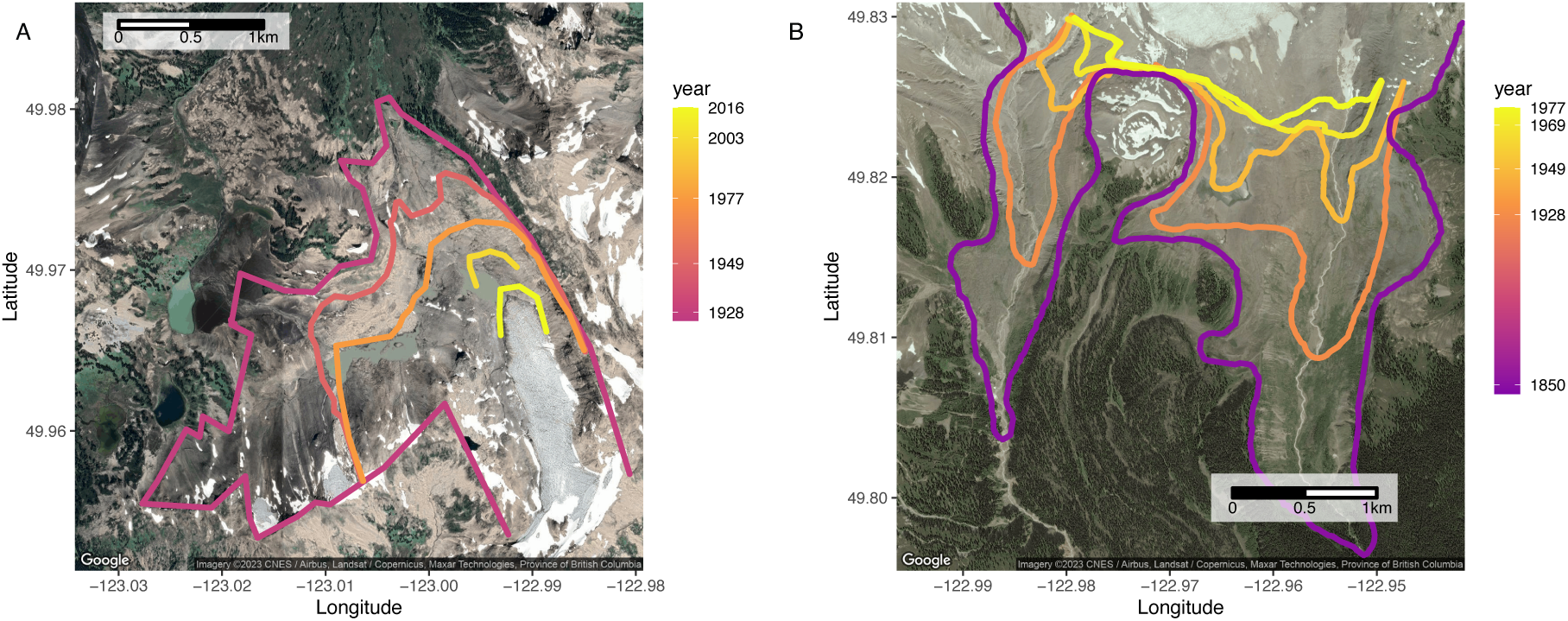
Mapped glacier recession pathways. **A**) Helm and **B**) Garibaldi/Lava from Koch et al. (2009). Habitat outside the earliest chronosequence zone is the same age for both forelands (< 1850). In the Helm glacier foreland, the glacier extent in 1928 advanced just beyond the LIA maximum (1850), giving the appearance of a potentially younger historical core habitat, however, habitat beyond the 1928 zone is of equivalent age to the Garibaldi/Lava historical core. For A), Helm, the glacier previously wrapped around Cinder Cone, with ice on the West (Cinder Flats) and East side (Alpine Meadows). For B), Garibaldi/Lava, the glacier wrapped around Opal Cone and down two river valleys (Garibaldi Glacier down Ring Creek to the west and Lava Glacier down Zig-Zag creek to the east.

**Figure S2:**
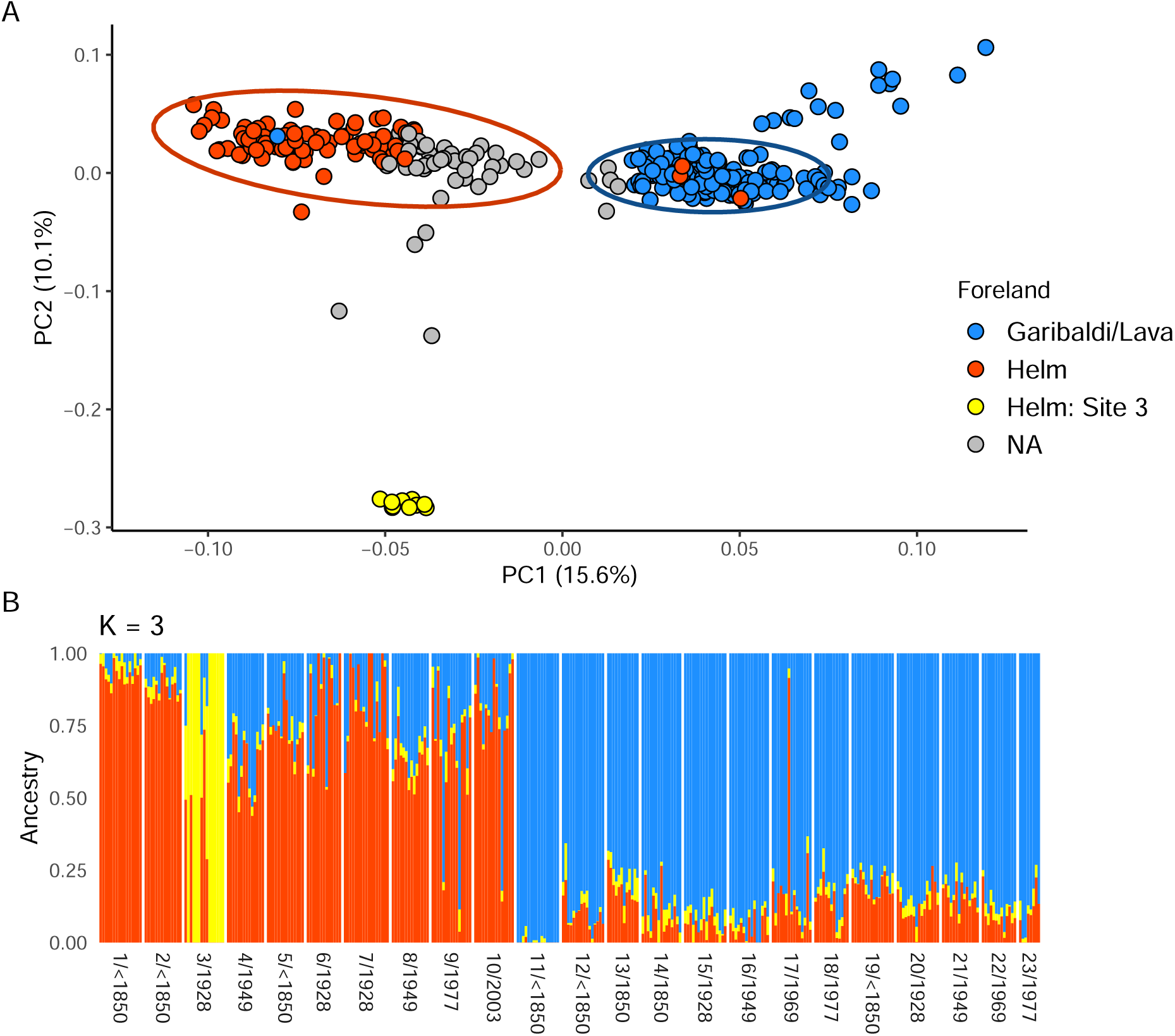
PCA and ADMIXTURE plot for two ancestry clusters (*K* = 3). A) Data points are coloured by by > 70% ancestry confidence for foreland clusters, grey points are individuals with ancestry assignment < 70% for each cluster. B) Structure plot from ADMIXTURE analysis with *K* = 3, identifying Site 3 (Helm glacier, located in chronosequnce zone 1928) coloured in yellow as generally genetically distinct.

**Figure S3:**
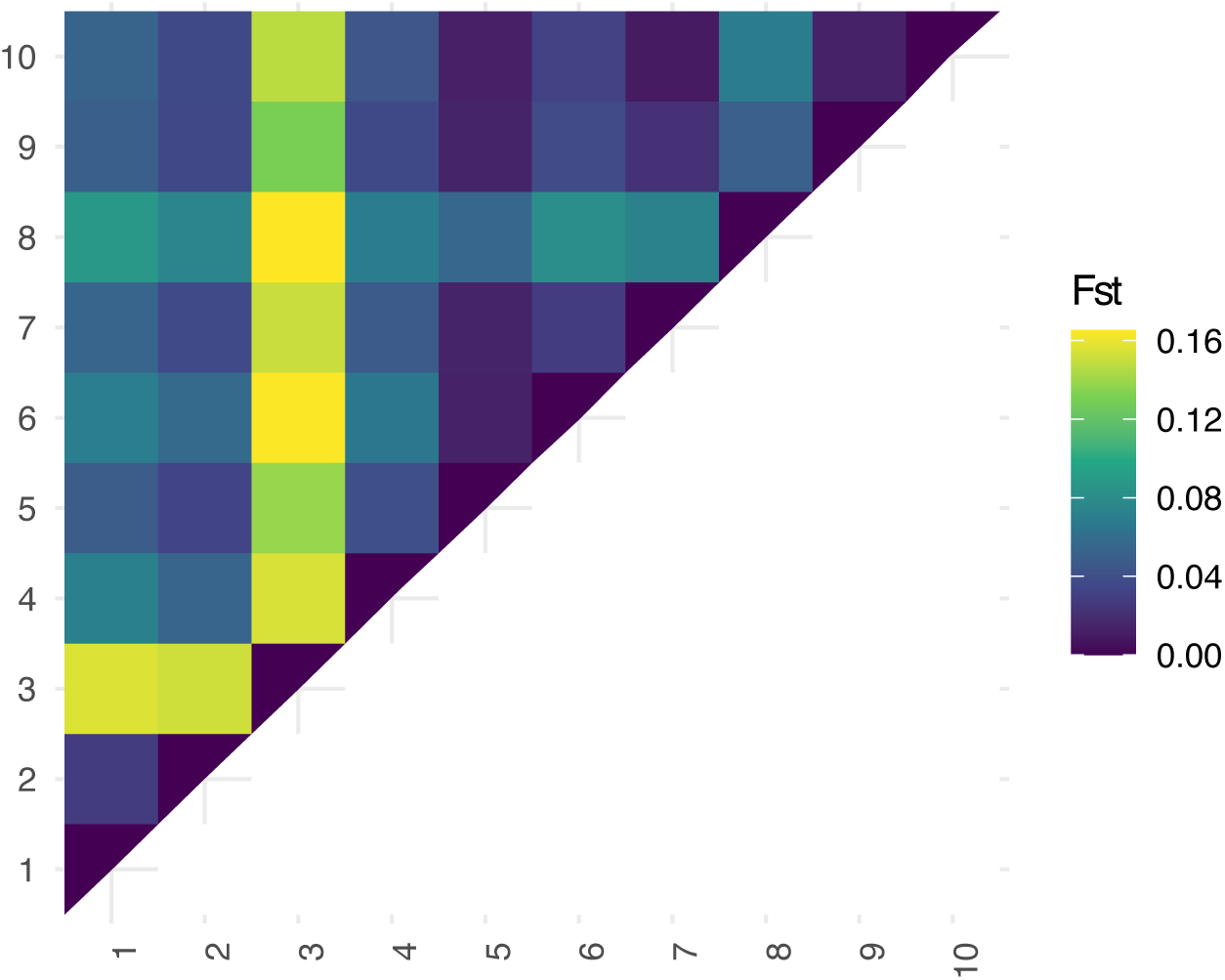
Heatmap of pairwise F_ST_ values for the Helm Glacier foreland. Note, pairwise comparisons with site 3 show relatively higher levels of genetic differentiation. Site numbers for rows and columns correspond to sampling locations and chronosequence year in Fig. 1 and Fig. 2B/S2B

**Figure S4:**
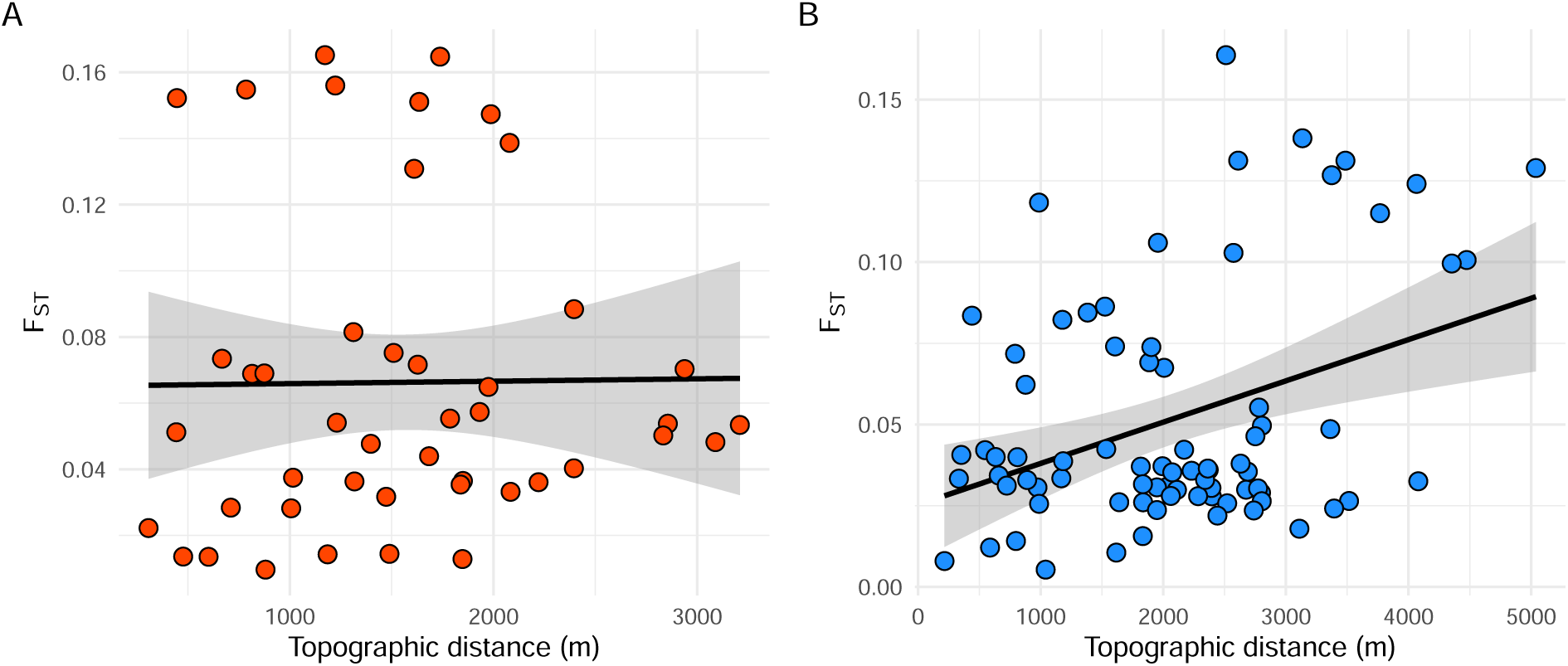
No evidence of restricted gene flow with topographic distance. Relationship between topographic distance among sampled sites within a foreland versus genetic differentiation (isolation-by-topographic-distance) (*F_ST_*) is non-significant in A) Helm Mantel statistic *r* : 0.117, *p* = 0.272) and B) Garibaldi/Lava forelands Mantel statistic *r* : 0.183, *p* = 0.141). Linear regression was used to fit lines to the scatterplots.

